# CDK8 inhibitors antagonize HIV-1 reactivation and promote provirus latency in T cells

**DOI:** 10.1101/2023.06.07.544021

**Authors:** Riley M. Horvath, Zabrina L. Brumme, Ivan Sadowski

**Affiliations:** Department of Biochemistry and Molecular Biology Molecular Epigenetics Group, LSI University of British Columbia, Vancouver, B.C., Canada; Faculty of Health Sciences, Simon Fraser University, Burnaby, B.C., Canada; British Columbia Centre for Excellence in HIV/AIDS, Vancouver, B.C., Canada

**Keywords:** CDK8, inhibitors, Senexin A, BRD6989, HIV-1, Latency, LTR, Transcription, mediator kinase, T cell signaling, block and lock

## Abstract

Latent HIV-1 provirus represents a barrier towards a cure for infection, but is dependent upon the host RNA Pol II machinery for expression. We find that inhibitors of the RNA Pol II mediator kinases CDK8/19, Senexin A and BRD6989, inhibit induction of HIV-1 expression in response to latency reversing agents and T cell signaling agonists. These inhibitors were found to impair recruitment of RNA Pol II to HIV-1 LTR. HIV-1 expression in response to several latency reversal agents was impaired upon disruption of *CDK8* by shRNA or gene knockout. However, the effects of CDK8 depletion did not entirely mimic CDK8/19 kinase inhibition suggesting that the mediator kinases are not functionally redundant. Furthermore, treatment of CD4^+^ PBMCs isolated from people living with HIV-1 and who are receiving ART with Senexin A inhibited induction of virus replication in response to T cell stimulation by PMA and ionomycin. These observations indicate that the mediator kinases CDK8 and CDK19, play a significant role for regulation of HIV-1 transcription, and that small molecule inhibitors of these enzymes may contribute to therapies designed to promote deep latency involving the durable suppression of provirus expression.

## Introduction

The ability of HIV-1 to persist within infected cells as an integrated provirus, even during suppressive antiretroviral therapy (ART), represents the main barrier to achieving ART-free HIV remission or cure^1 2^. Various strategies to inactivate or eliminate latent viral reservoirs have been proposed, many of which involve approaches to modulate proviral expression^1 3^. Like most cellular genes, HIV-1 provirus within latently infected cells produce stochastic transcripts through transcriptional noise^4 5^, and there is further speculation that such transcripts may contribute to HIV reservoir maintenance in individuals receiving ART^4^. Consequently, it has been proposed that therapies capable of durably suppressing basal stochastic expression might allow treatment to be removed without the risk of viral rebound. This rationale is the basis for the HIV remission strategy known as "block and lock", where intervention would be applied to durably suppress basal and stochastic provirus expression^6 7 8^. Much focus towards this objective has centered on the viral Tat protein, which binds the nascent TAR RNA and recruits P-TEFb, containing CDK9 and Cyclin T1, which phosphorylates proteins at the core promoter to encourage transcriptional elongation by RNAP II ^9^. Inhibitors of Tat block the positive feedback effect caused by this factor in cells producing basal provirus transcripts^10 11^. Additional potential block and lock targets under current investigation include factors required for transcriptional initiation or elongation, including inhibitors of CDK9^12^.

Expression of chromosomally integrated HIV-1 provirus in T cells is regulated by multiple signaling pathways that are activated in response to stimulation by cytokines, innate immune responses, and engagement of the T cell receptor with antigen presenting dendritic cells. Responses to these signals are controlled by sequence-specific transcriptional activator proteins bound to the 5’ LTR enhancer region, including NF-κB, NFAT, GABP/ Ets, STAT1/3, AP1, TCF-1/ Lef, and TFII-I-USF1/2 (RBF-2)^13 14^. These factors stimulate transcription from the HIV-1 LTR promoter through interactions that cause recruitment of general transcription factors (GTFs) for RNA Polymerase II, in addition to co-activator complexes, including the mediator^15 16^. Several factors bound to the LTR, including TFII-I/ USF1/2 (RBF-2) also promote transcription elongation by recruiting CDK9/ P-TEFb through interaction with the co-activator TRIM24^17 18^.

The RNA Polymerase II mediator kinase submodule is comprised of CDK8, or its paralog CDK19, cyclin C, and the regulatory factors Med12 and Med13. This submodule transiently associates with the core mediator, is recruited to promoters by transcriptional activators^19^, and acts to modulate transcription by phosphorylating sequence specific activators as well as GTFs^20^. Defined GTF substrates for CDK8/19 include the RNA Pol II B220 C-terminal domain (CTD) and mediator subunits^21 22 19^. Phosphorylation of transcriptional activators by CDK8/19 can have positive or negative effects on transcription, depending on the functional effect of the modification^19^. CDK8-dependent phosphorylation of β-catenin TCF/LEF^23^, NF-κB p65^24^, and STAT1/3^25^, factors that regulate expression of HIV-1 provirus in response to T cell signaling, enhances transcription. In contrast, the notch intracellular domain (NICD) is negatively regulated by CDK8 phosphorylation, which promotes its degradation^26^.

Consistent with observations that CDK8 phosphorylation has both positive and negative effects on transcription factors, alterations in the *CDK8* and cyclin C (*CCNC*) genes have been implicated as both oncogenes and tumor suppressors. *CDK8* is overexpressed in a variety of cancers, including breast and colorectal carcinomas, and malignant melanomas, where expression is associated with tumor progression^23^. In contrast, *CDK8* overexpression produces tumor suppressive effects of cancers promoted by Notch or EGFR signaling^27^. The significance for alterations in mediator kinase components and cancer progression has spurned development of specific CDK8/19 inhibitors^28 29 30^, several of which are in clinical trials for ER-positive breast cancers and acute myeloid leukemia^31^. Despite that HIV-1 expression is regulated by at least three factors whose activity is controlled by CDK8, TCF/LEF, NF-κB, and STAT1/3, the role of CDK8/19 and the kinase module for regulation of HIV-1 provirus expression and response to T cell signaling has not been characterized.

In this report, we examine the effect of CDK8/19 inhibitors and *CDK8* knockouts on expression of HIV-1 provirus. We find that Senexin A and BRD6989, structurally distinct chemical inhibitors specific for CDK8/19 kinase activity, impair reactivation of HIV-1 provirus in cell line models of latency, and encourage establishment of immediate latency in newly infected T cells. Inhibition of CDK8/19 impairs recruitment of RNA Pol II to the LTR promoter. Furthermore, CRISPR-mediated knockout of the *CDK8* gene impairs the expression and activation of HIV-1 in response to several latency reversal agents. Interestingly however, disruption of *CDK8* does not entirely mimic the effect of CDK8/19 kinase inhibition, indicating that CDK8 and CDK19 do not function redundantly for the activation of proviral transcription. Collectively, our observations indicate that CDK8/19 inhibitors, including those currently in clinical trials for treatment of various cancers, may prove useful for therapies to eliminate latently infected cells by suppressing HIV-1 expression.

## Results

### Chemical inhibition of CDK8/19 kinase activity suppresses HIV-1 expression

We examined the effect of small molecule inhibitors of CDK8/19 kinase activity on HIV-1 expression by treating the previously characterized JLat10.6^32^ and mHIV-Luciferase Jurkat^33^ human CD4^+^ T cell lines with two structurally unrelated CDK8/19 inhibitors, Senexin A and BRD6989 (Fig. 1*A*)^34 35^. JLat10.6 Jurkat cells possess a full-length HIV-1 provirus with GFP expressed instead of *Nef* from the 5’ LTR^71^, while Jurkat Tat mHIV-Luciferase cells express luciferase as a fusion with gag from the 5’ HIV-1 LTR, but no other HIV-1 proteins (Fig. 1*B*)^33 78^. We did not observe an effect on basal GFP expression in JLat10.6 cells upon treatment with these inhibitors, but induction of GFP expression in cells treated with the PKC agonist PMA was inhibited in a concentration-dependent manner (Fig. 1*C*, 1*D*). A similar effect was seen with mHIV-Luciferase cells, although with this reporter line we also recorded a significant reduction in basal luciferase expression upon treatment with Senexin A but not with BRD6989 (Fig. 1*E*, 1*F*). The inhibitory effect of the CDK8/19 inhibitors on HIV-1 expression was not due to toxicity, as concentrations as high as 100 μM had only minor effects on cell viability (Fig. S1*A*, S1*B*). Additionally, we examined effect of the CDK8/19 inhibitors on expression of HIV-1 provirus in the ACH2 CEM CD4^+^ T cell and U1 monocyte models of latency, where we likewise observe inhibition of basal and PMA-induced expression of viral mRNA (Fig. 2*A*, 2*B*) and no toxicity at concentrations where inhibition was detected (Fig. S1*C*, S1*D*). These observations indicate that chemical inhibition of CDK8/19 kinase activity suppresses HIV-1 expression in several cell line models of provirus latency.

**Figure 1.**
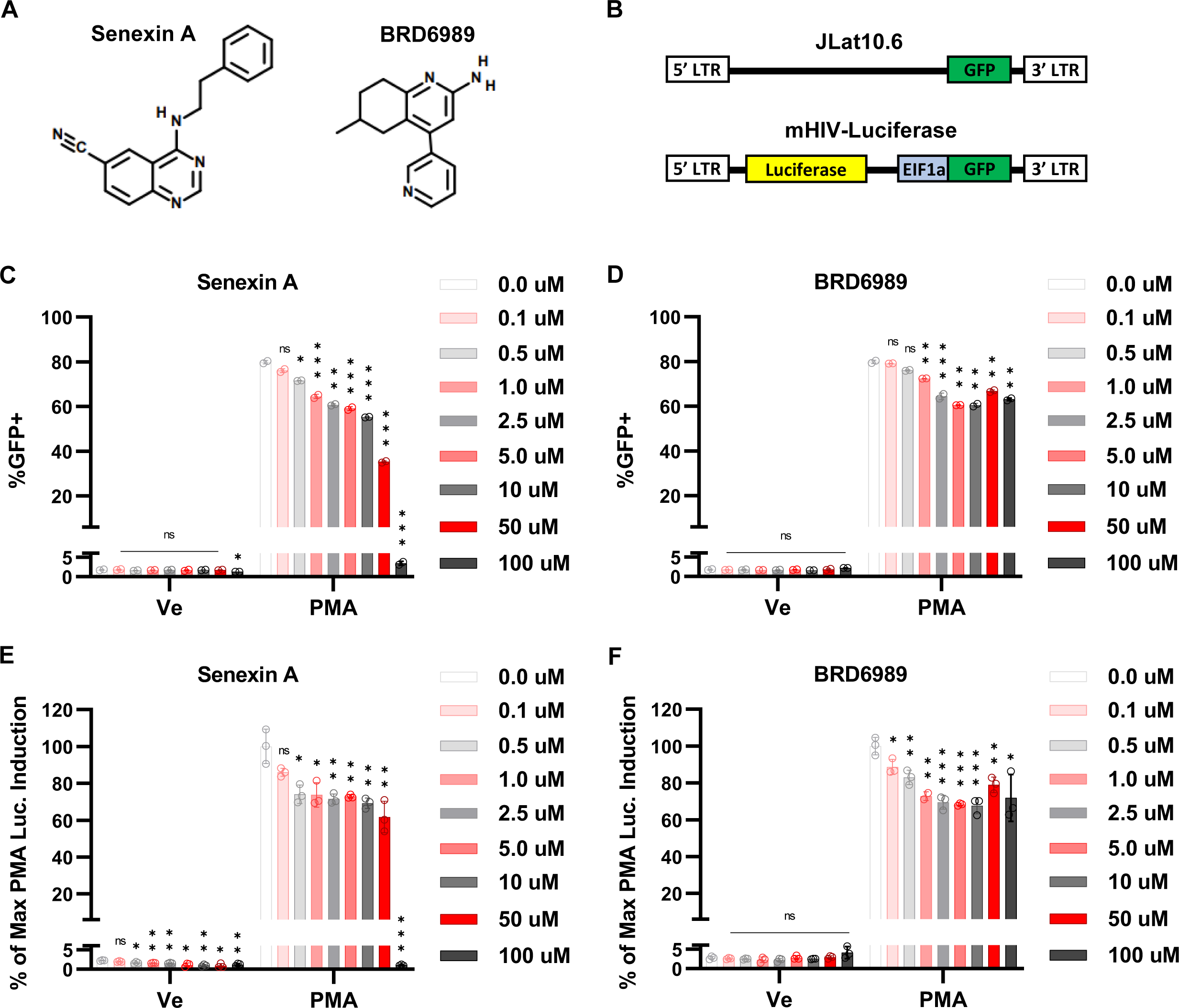
CDK8/19 inhibitors impair reactivation of HIV-1 provirus. **A:** Structure of CDK8/19 inhibitors Senexin A and BRD6989. **B:** Schematic representation of integrated reporter virus in the JLat10.6 (top) and mHIV-luciferase (bottom) Jurkat cell lines. The JLat10.6 reporter represents the full HIV-1 genome with GFP inserted into *nef*, while mHIV-luciferase cells express luciferase as a fusion with gag from the 5’ LTR, with a provirus that is deleted of all other viral encoded proteins. **C, D:** JLat10.6 cells were pre-treated for 1 hr with the indicated concentration of Senexin A (C) or BRD6989 (D). Subsequently, 10 nM PMA was added and flow cytometry was performed following 20 hrs of incubation (*n* = 2, mean ± SD). **E, F:** mHIV-Luciferase cells were pre-treated for 1 hr with the indicated concentration of Senexin A (E) or BRD6989 (F). Following pre-treatment, 10 nM PMA was added, and luciferase assays were performed after 4 hrs. Results are normalized to the maximum luciferase expression recorded, i.e., PMA induced mHIV-Luciferase cells that were not treated with any amount of Senexin A or BRD6989 (*n* = 3, mean ± SD).

**Figure 2.**
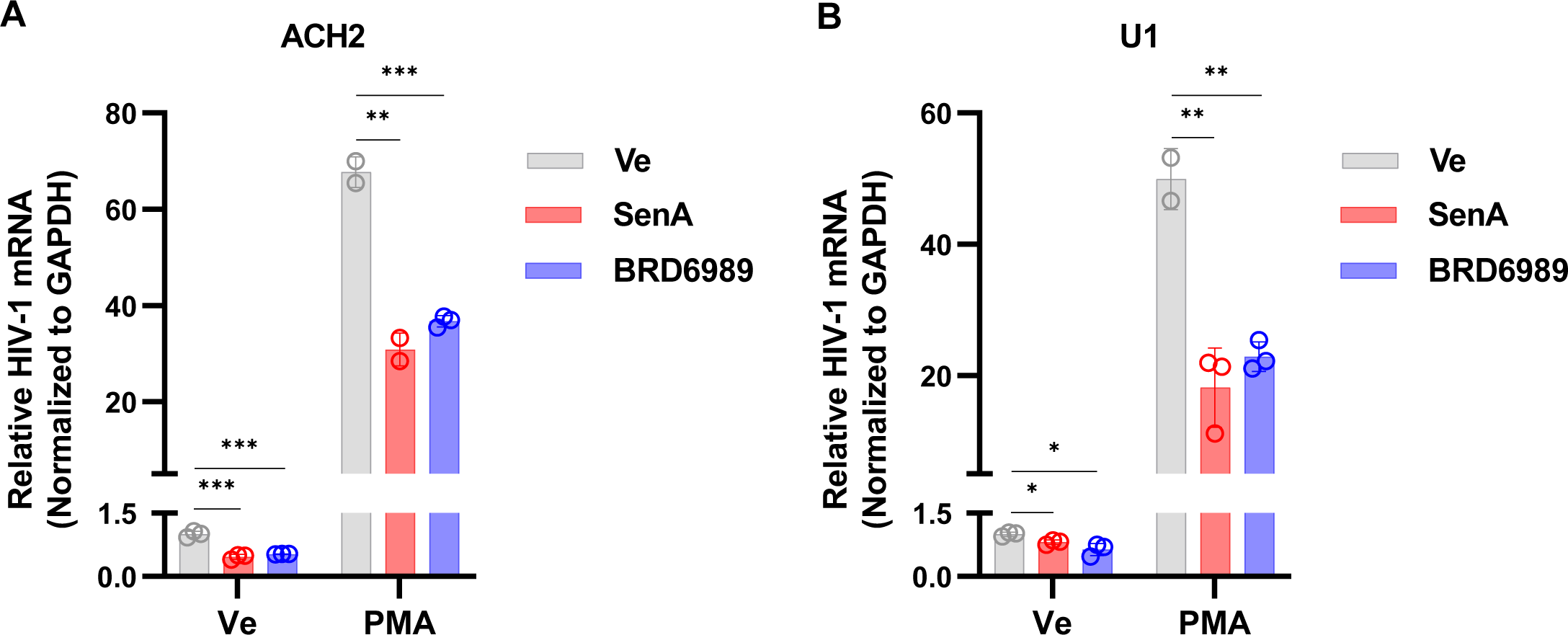
CDK8/19 inhibitors prevent reactivation of HIV-1 in cell models of HIV-1 latency. **A, B:** ACH2 (A) or promonocytic U1 cells (B), were incubated with a DMSO vehicle control (Ve), 10 μM Senexin A, or 10 μM BRD6989 for 1 hr. Afterwards, DMSO (Ve) or 10 nM PMA was added, and cells were incubated for 20 hrs prior to extraction of intracellular RNA. Expression of HIV-1 mRNA was measured by performing RT-PCR with oligos that detect multiply spliced Tat-Rev transcripts and normalizing to *GAPDH* (*n* = 2 - 3, mean ± SD).

### CDK8/19 antagonizes HIV-1 reactivation by latency reversing agents

We examined the effect of CDK8/19 inhibitors on stimulation of HIV-1 expression by additional signaling agonists. Treatment with a combination of the phorbol ester PMA and ionomycin mimics T cell activation by stimulation of the MAPK, PKC and calcineurin signaling pathways downstream of the T cell receptor^36 37^. PMA stimulates PKC, activating the MAPK and IKK-IκB-NFκB pathways, while ionomycin causes release of intracellular calcium resulting in activation of NFAT through calcineurin mediated dephosphorylation^14^. We found that HIV-1 expression in both the JLat10.6 and mHIV-Luciferase cell lines treated with PMA, alone or in combination with ionomycin, is inhibited by Senexin A and BRD6989 (Fig. 3*A*, 3*B*). Similarly, the CDK8/19 inhibitors also significantly impair HIV-1 activation in response to Ingenol 3-angelate (PEP005) (Fig. 3*A*, 3*B*), a latency reversal agent (LRA) which activates NFκB^38 39^.

**Figure 3.**
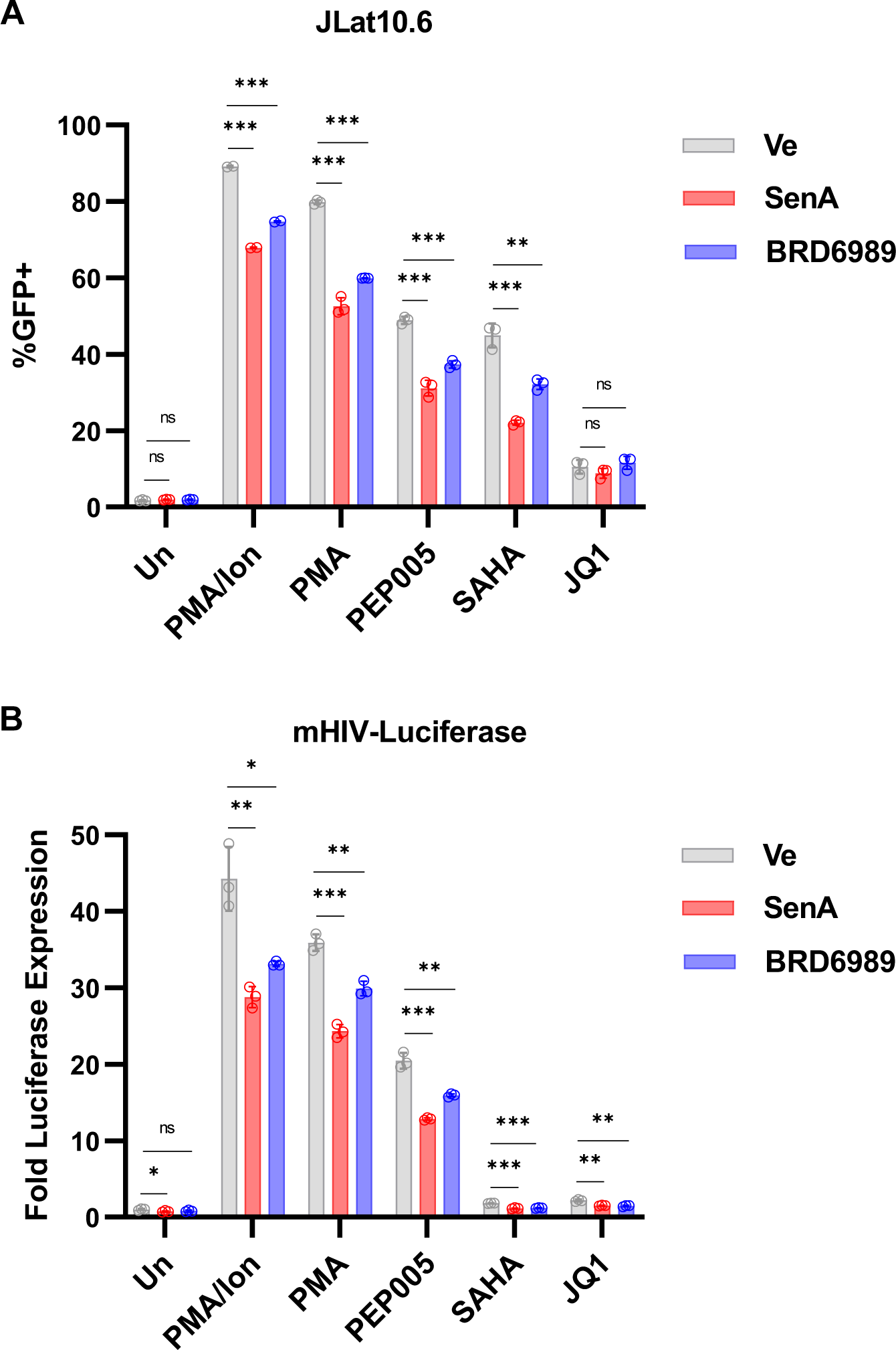
Inhibition of CDK8/19 impairs reactivation of HIV-1 by latency reversing agents. **A, B**: JLat10.6 (A) or mHIV-Luciferase (B) cells were treated with a DMSO vehicle control (Ve), 10 μM Senexin A, or 10 μM BRD6989 for 1 hr. Next, cells were left untreated (Un) or treated with 10 nM PMA/ 1 μM ionomycin, 10 nM PMA, 10 nM PEP005, 10 μM SAHA, or 10 μM JQ1. GFP expression of JLat10.6 cells was determined by flow cytometry following 20 hrs of incubation (*n* = 2 - 3, mean ± SD). mHIV-Luciferase cells were treated as in (A), but LTR expression was determined following 4 hrs treatment by luciferase assay (*n* = 3, mean ± SD).

Latent HIV-1 provirus can also be reactivated by treatments which affect chromatin modifications, or association of epigenetic modifiers with the LTR in the absence of signaling agonists^40^. The histone deacetylase inhibitor (HDACi) suberanilohydroxamic acid (SAHA) for example, as well as the BRD4 inhibitor JQ1, are well characterized latency reversing agents (LRAs)^41 39^. SAHA induces expression of HIV-1 to a comparable level as PEP005 in the JLat10.6 cell line, and we found that this effect was also inhibited by Senexin A and BRD6989 (Fig. 3*A*). The effect of SAHA on HIV-1 expression in the mHIV-Luciferase line was not as pronounced, and here we observed only a minor but significant effect of the CDK8 inhibitors (Fig. 3*B*). JQ1 caused significantly lower levels of reactivation in both reporter cell lines, which was only significantly affected by the CDK8/19 inhibitors in the mHIV-Luciferase cell line (Fig. 3*A*, 3*B*). These results demonstrate that CDK8/19 kinase activity is required for the most robust reactivation of HIV-1 expression in response to a variety of LRAs.

### Effect of CDK8/19 inhibition on cell growth

Previously, we investigated the effect of Senexin A and BRD6989 on HIV-1 expression by employing temporally acute treatments. To further examine the cellular impact of CDK8/19 inhibition, we assessed the growth of Jurkat T cells exposed to a range of drug concentrations for four days (Fig. 4). On day 3, 10 μM of Senexin A was found to slightly impair cell expansion while 50 μM of the compound elicited cell cycle arrest (Fig. 4*A*). In contrast, cell cycle progression was less affected by BRD6989 with concentrations as high as 50 μM having minor observable effects on cell division (Fig. 4*B*). Importantly, no effect on cellular viability throughout the four days examined was observed for any of the employed Senexin A or BRD6989 concentrations (Fig. 4*C*, 4*D*).

**Figure 4.**
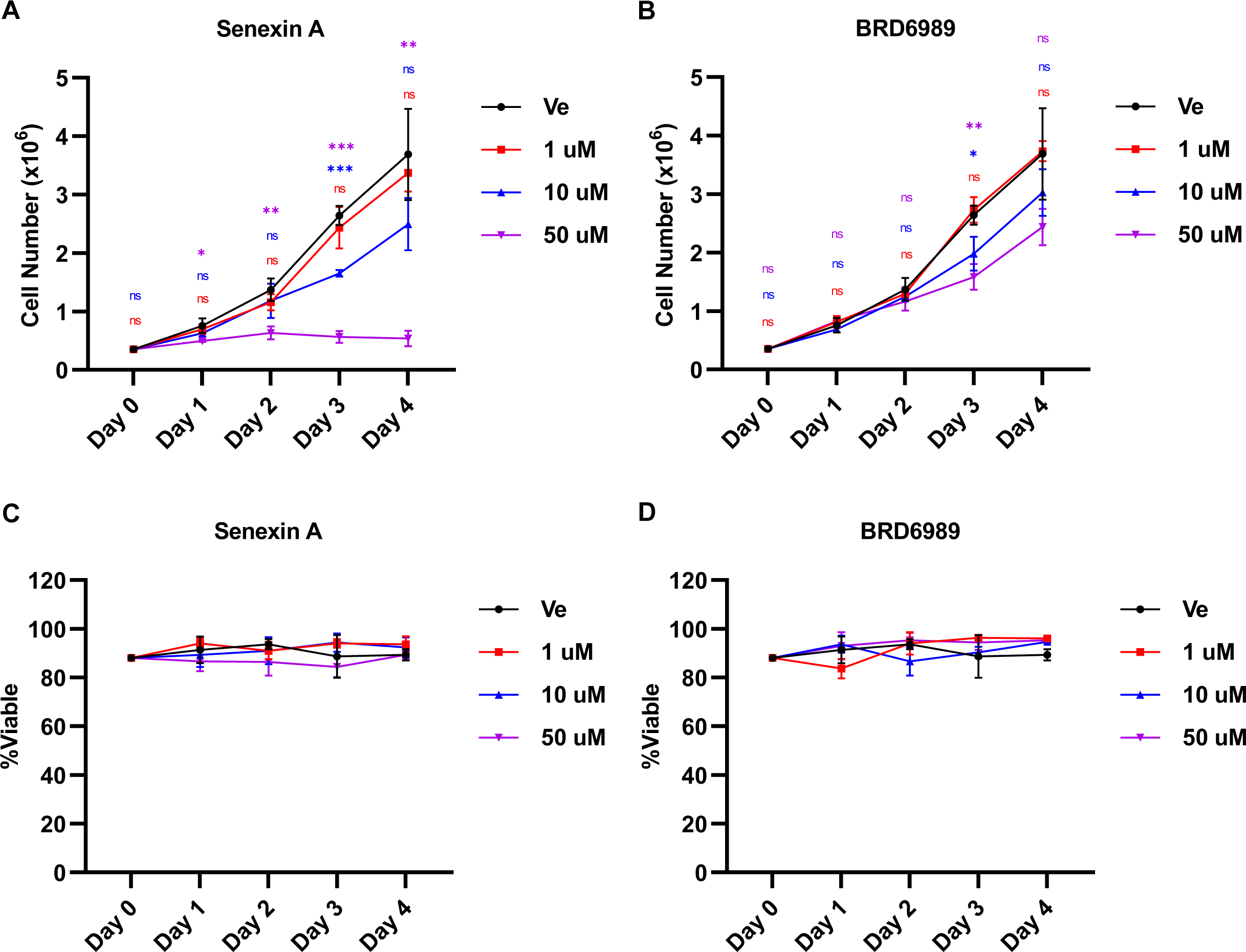
Effect of Senexin A and BRD6989 on cellular growth and viability. **A, B:** Jurkat T cells were incubated with the indicated concentration of Senexin A (A) or BRD6989 (B) or a vehicle control (Ve, DMSO) for four days. Cells were counted on the indicated day for the given treatment (*n* = 3, mean ± SD). **B, C:** Jurkat T cells treated as in A, B were assessed for viability (*n* = 3, mean ± SD).

### Inhibition of CDK8/19 encourages establishment of HIV-1 latency

Having determined concentrations of Senexin A and BRD6989 that have no observed toxicity and minimal impact on cell cycle progression, we next examined the role of CDK8/19 for establishment of immediate latent infection in Jurkat T cells using the replication incompetent Red-Green-HIV-1 (RGH) virus. RGH is a dual reporter virus where GFP is expressed from the 5’ LTR and mCherry from an internal constitutive PGK promoter, thus allowing latently infected (mCherry+/GFP-) cells to be discriminated from productively infected (mCherry+/GFP+) cells as early as 24 hours post infection by flow cytometry (Fig. 5*A*, 5*B*)^42 43^. Consistent with previous observations^44 43^, the proportion of productively infected cells peaks at 4 days post infection, and then declines as the provirus succumbs to silencing (Fig. 5*C*, Ve)^44^. In contrast, infection of cells treated with Senexin A or BRD6989 over the course of the analysis produced a significantly lower proportion of productively infected cells, with the largest discrepancy occurring at 7 days post-infection (Fig. 5*B*, 5*C*). Similar results were observed with infection of the SupT1 human T cell line (Fig. S2*A,* S2*B*), and with an additional dual HIV-1 reporter virus (Fig. S2*C*, S2*D*)^45^. These results indicate that inhibition of CDK8/19 kinase activity in newly infected cells encourages production of immediate latent provirus.

**Figure 5.**
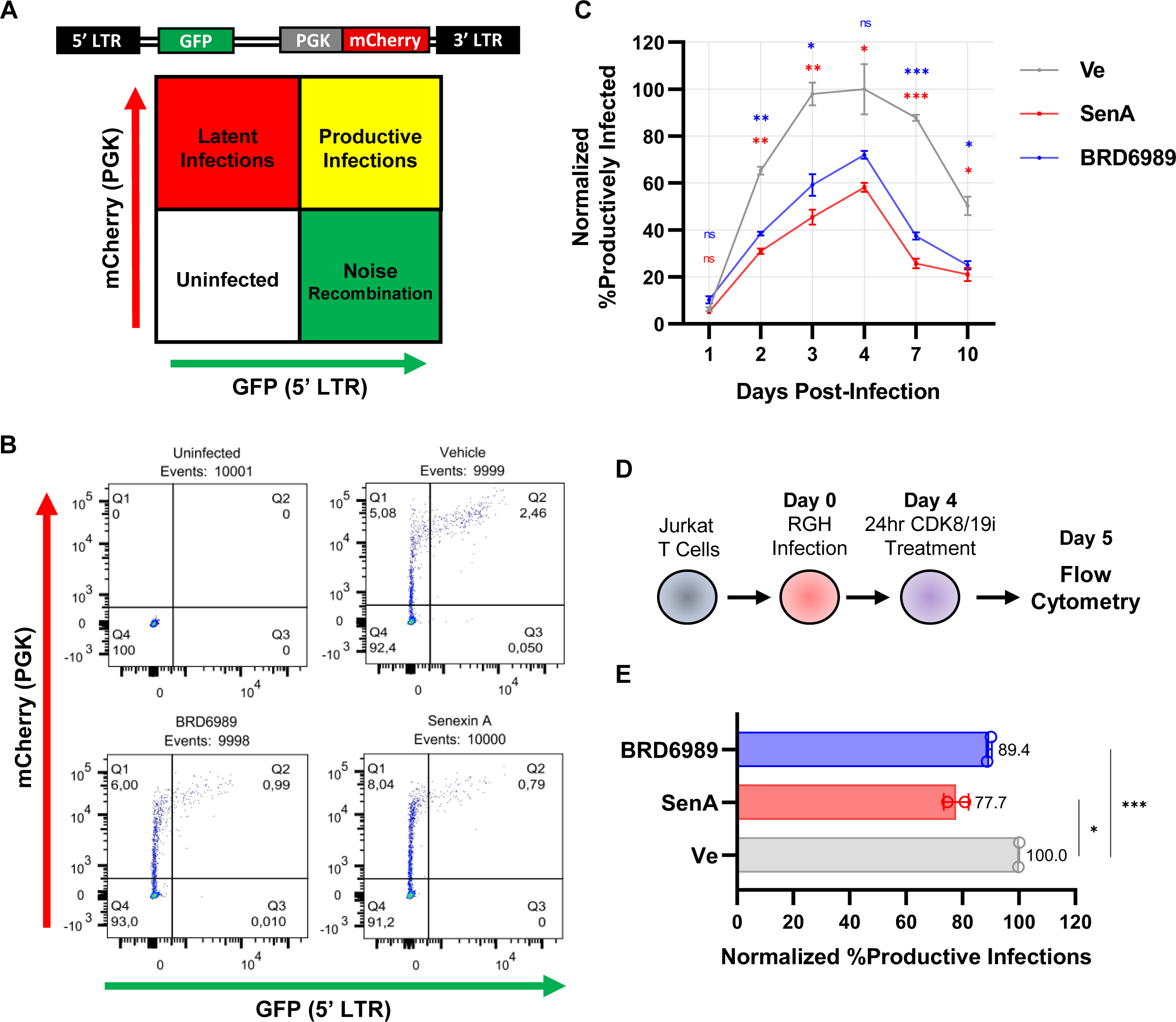
Inhibition of CDK8/19 promotes establishment of latency in newly infected T cells. **A:** Flow cytometric analysis of cells infected with the single-round, Red-Green-HIV-1 (RGH) dual reporter virus produces a scatter plot with distinct populations. Bottom left contains the uninfected (fluorescent negative) cells, top left is the population of latently infected cells (mCherry+/GFP-), top right displays the productive infections (mCherry+/GFP+), while bottom right contains noise generated from viral rearrangements (mCherry-/GFP+). **B:** Representative flow cytometry scatter plots of Jurkat T cells that are uninfected or are 7 days post RGH infection while being treated with a vehicle control (DMSO), 10 μM Senexin A, or 10 μM BRD6989. **C:** Untreated (Ve, DMSO) or Jurkat T cells treated with 10 μM Senexin A or 10 μM BRD6989 were infected with RGH, and the percentage of productively infected cells was determined at the indicated times post-infection by flow cytometry. Values are normalized to the ratio of productive infections of the vehicle control (Ve) (*n* = 2, mean ± SD). **D:** The effect of treatment with CDK8/19 inhibitors post-infection was determined by the addition of 10 μM Senexin A or 10 μM BRD6989 to Jurkat cells 4 days post-infection with RGH, and the proportion of cells with productive infections was determined 20 hours later by flow cytometry. **E:** The proportion of productively infected cells treated 4 days post-infection with 10 μM Senexin A or 10 μM BRD6989, was determined by flow cytometry. Values are normalized to the ratio of productive infections of the vehicle control (Ve) (*n* = 2, mean ± SD).

Additionally, we examined whether treatment of cells with the CDK8/19 inhibitors post-infection affected the proportion of latent or productively infected cells. For this, we treated cells with the CDK8/19 inhibitors 4 days post-infection and measured the proportion of latently and productively infected cells 24 hours later (Fig. 5*D*). In this experiment we found that Senexin A caused a ∼20% reduction in productive infections, while BRD6989 caused a more modest ∼10% decrease as compared to cells treated with a vehicle control (Fig. 5*E*). This result indicates that effects of the CDK8/19 inhibitors are not temporally restricted to immediate events in HIV-1 infection, including entry and formation of integrated provirus, but rather must inhibit expression of the virus post-integration, which is consistent with the effect of these compounds on reactivation of provirus reporter by latency reversing agents.

### CDK8/19 inhibitors encourage HIV-1 latency in primary CD4^+^ T cells ex vivo

Having observed that CDK8/19 inhibition suppresses HIV-1 expression in immortalized human T cell lines, we next examined the effect of Senexin A and BRD6989 on HIV-1 latency in primary CD4^+^ T cells. For this, we treated CD4^+^ T cells from normal donors with the CDK8/19 inhibitors or a vehicle control prior to infection with the replication incompetent RGH dual-reporter virus. Proviral activity was assessed by flow cytometry 3 days post-infection (Fig. 6*A*). Consistent with the results described above with immortalized T cell lines, we found that Senexin A and BRD6989 cause a significant decrease in the proportion of productively infected cells, represented by expression of both mCherry and GFP as compared to mCherry alone (Fig. 6*B*), where Senexin A inhibited to a greater effect than BRD6989 (Fig. 6*C*, 6*D*). Furthermore, we found that proviruses that managed to establish productive infections in the presence of Senexin A or BRD6989 displayed lower levels of transcriptional activity than their untreated counterparts (Fig. 6*D*, Productive Inf.). Notably, the CDK8/19 inhibitors did not affect the viability (Fig. 6*E*) and only had a slight impact on division (Fig. 6*F*) of the primary CD4^+^ T cells over the course of the treatment, similar to the results obtained from Jurkat T cells (Fig. 4). Finally, primary CD4^+^ T cells treated with either CDK8/19 inhibitor showed no difference in susceptibility to HIV-1 infection (Fig. 6*G*). Collectively, these results indicate that Senexin A and BRD6989 inhibit HIV-1 expression in primary T cells, an effect which encourages the formation of latency.

**Figure 6.**
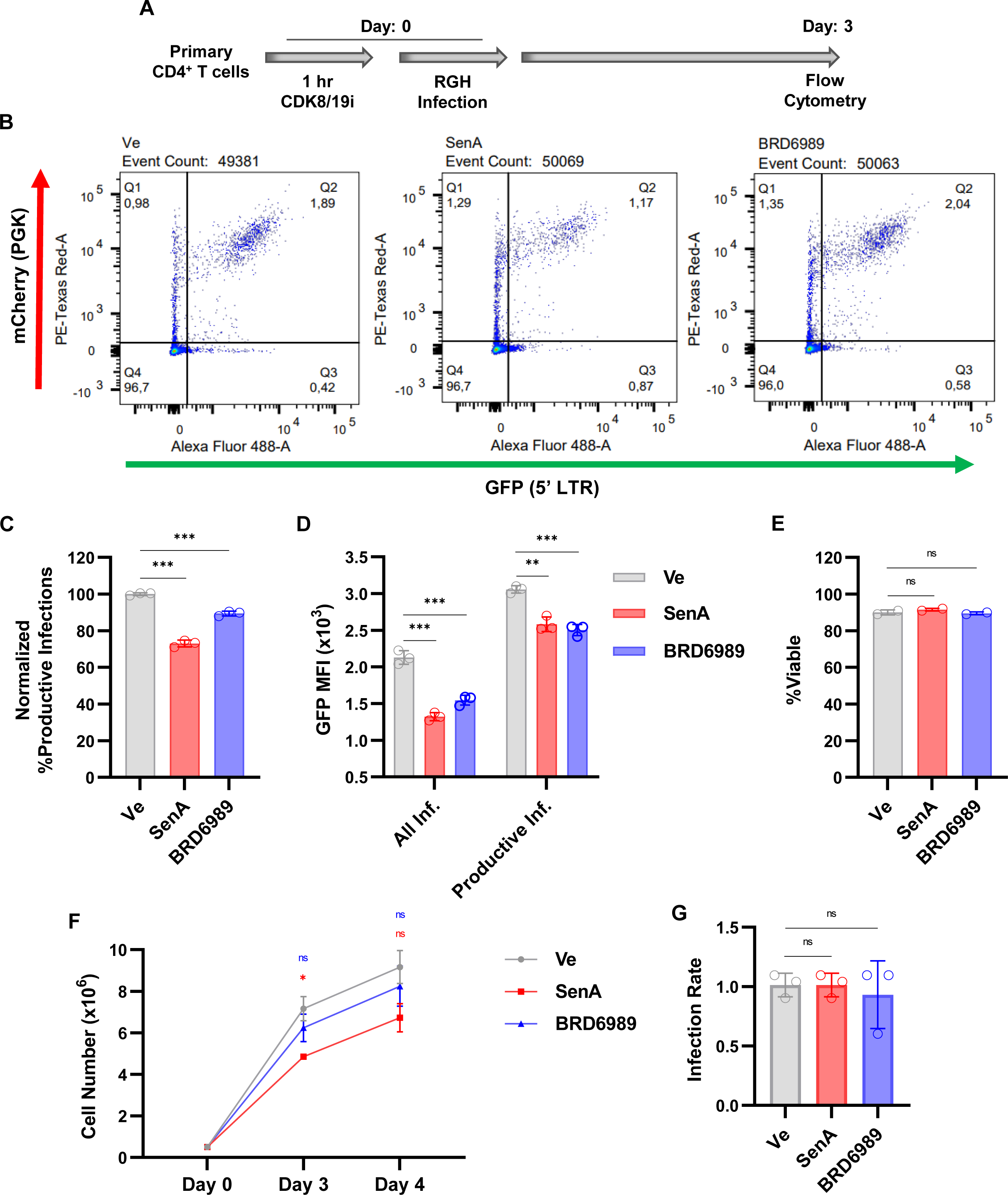
HIV-1 latency is regulated by CDK8/19 kinase activity in primary CD4^+^ T cells. **A:** Following a 1 hr pre-treatment with a vehicle control (Ve, DMSO), 10 μM Senexin A, or 10 μM BRD6989, CD4^+^ PBMC T cells were infected with RGH. Flow cytometry was performed after three days of treatment. **B:** Representative flow cytometry scatter plots of primary CD4^+^ T cells infected with RGH. GFP expression is indicated on the x-axis and mCherry on the y-axis. Q1 is the population of cells that is latently infected (mCherry+/GFP-), Q2 displays productive infections (mCherry+/GFP+), Q3 depicts noise generated from viral rearrangements (mCherry-/GFP+), and Q4 displays the uninfected population (mCherry-/GFP-). **C:** RGH infected CD4^+^ T cells were examined by flow cytometry 3 days post-infection. Values are normalized to the ratio of productive infections of the vehicle control (Ve) (*n* = 3, mean ± SD). **D:** As in (C) but the Mean Fluorescence Intensity (MFI) of GFP expression is assessed. GFP MFI was evaluated within the entire population of infected cells (All Infections, mCherry+/GFP- and mCherry+/GFP+) or within the population of cells that harbored productive infections (Productive, mCherry+/GFP+) (*n* = 3, mean ± SD). **E:** Viability of cells treated as in (A) (*n* = 2, mean ± SD). **F:** Primary CD4^+^ T cells treated as in (B) were counted on the indicated day post-infection (*n* = 2, mean ± SD). **G:** The infection rate was determined for primary T cells as treated in (B). Values are normalized to the vehicle control (Ve) (*n* = 3, mean ± SD).

### Senexin A and BRD6989 do not inhibit HIV-1 Tat, and block HIV reactivation in a cell-specific manner

Originally isolated from the murine sponge *Corticium simplex* ^46^, the steroidal alkaloid cortistatin A (CA) was found to inhibit CDK8/19 kinase activity^47 48^. Because of its limited availability from nature, analogs of CA were developed, including didehydro-Cortistatin A (dCA)^49 50^. Subsequently, dCA was found to suppress HIV-1 expression by a mechanism proposed to involve inhibition of Tat, the HIV-1 transactivator of transcription, independently from effects involving inhibition of CDK8/19 activity^51 11 52 12^. Given these reported effects of dCA on HIV-1 Tat, we examined whether the CDK8/19 inhibitors Senexin A and BRD6989 had a similar effect on Tat function. To examine this, we transfected HEK293T cells with an HIV-1 LTR reporter plasmid where GFP is expressed from an IRES, alone or in combination with viral Tat protein (Fig. 7*A*). As expected, transfection of HEK293T cells with the reporter co-expressing Tat resulted in substantially elevated GFP expression compared to the reporter lacking Tat (Fig. 7*B*). Interestingly however, neither of the CDK8/19 kinase inhibitors had an effect on expression of GFP from either of these reporters (Fig. 7*B*), indicating that these compounds likely do not affect Tat function, unlike the reported effect of dCA^12 52^. Consistent with this observation, we note that HIV-1 provirus in the ACH2 cell line possesses a point mutation in TAR rendering it defective to Tat transactivation^53^, while the provirus in U1 cells expresses defective Tat protein^54^. As shown above (Fig. 2), Senexin A and BRD6989 both inhibit reactivation of HIV-1 expression in these lines. Furthermore, Senexin A and BRD6989 are not structurally related to dCA^34 35^, and these results indicate they must have distinct mechanisms of action.

**Figure 7.**
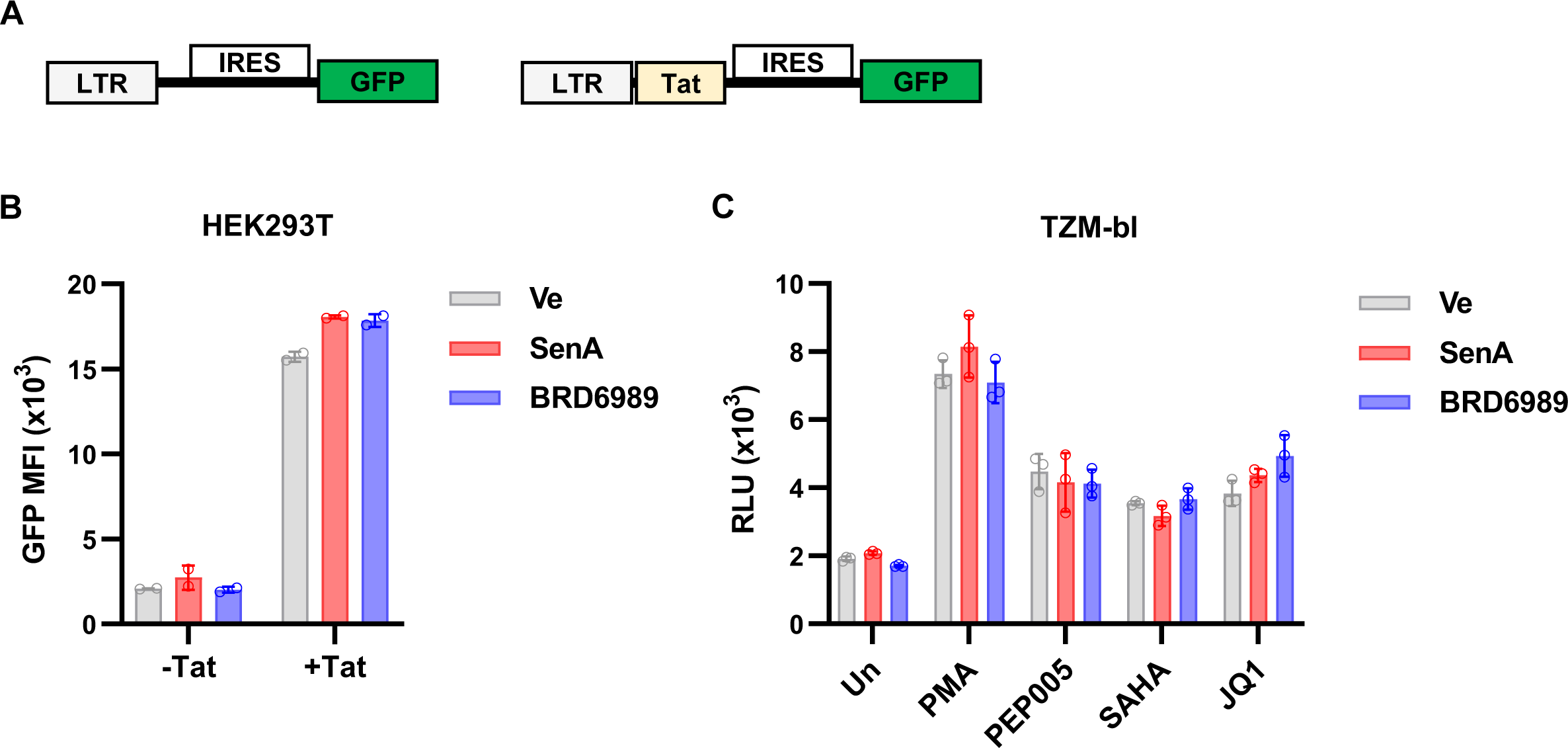
CDK8/19 inhibitors do not affect HIV-1 transactivation by Tat. **A:** Schematic representation of HIV-1 LAI derived LTR-IRES-GFP (-Tat) and LTR-Tat-IRES-GFP (+Tat) reporter constructs. **B:** HEK293T cells were transfected with the GFP reporters; one day post-transfection, cells were either left untreated (Ve) or treated with 10 μM Senexin A or 10 μM BRD6989. Flow cytometry was performed 20 hr later with the Mean Fluorescence Intensity (MFI) of the GFP+ population being measured (*n* = 2, mean ± SD). **C:** TZM-bl Hela-derived cells possessing an integrated LTR-luciferase reporter were pre-treated for 1 hr with DMSO vehicle control (Un), 10 μM Senexin A, or 10 μM BRD6989. The pre-treated cells were then stimulated with 10 nM PMA, 10 nM PEP005, 10 μM SAHA, or 10 μM JQ1. Luciferase expression was measured following 4 hrs of treatment and is displayed as Relative Light Units (RLU) (*n* = 3, mean ± SD).

Considering that the CDK8/19 inhibitors antagonize HIV-1 expression in Jurkat T cells but did not affect basal or Tat-activated expression of the LTR-IRES-GFP reporter in HEK293T cells (Fig. 7*B*), we wondered whether the effect of these compounds on HIV-1 expression may be cell type-specific. To further examine this, we used the HeLa-derived TZM-bl cell line which bears an integrated HIV-1 LTR-β-Gal-Luciferse reporter^55^. We observed induction of luciferase expression in this line in response to PMA, or the LRAs PEP005, SAHA and JQ1. However, treatment with Senexin A or BRD6989 had no effect on this response (Fig. 7*C*). These observations indicate that effects of CDK8/19 kinase activity on HIV-1 expression may be specific to cells of leukocyte lineage. Relating to this, we note that it was previously reported that knockdown of CDK8 and/or CDK19 does not affect HIV-1 infection in HeLa cells, where dCA has an inhibitory effect^12^. This supports a view that dCA must have additional inhibitory target(s) apart from the CDK8/19 kinase such as Tat, and that the effect of mediator kinase for HIV-1 expression may be unique to T cells.

### Inhibition of CDK8/19 inhibits recruitment of RNAPII to the LTR

Regulation of transcription from the HIV-1 LTR, like most cellular genes, involves the recruitment of RNA Pol II and histone modifying complexes to the promoter as well as regulation of transcriptional elongation and mRNA processing^56 1^. To examine which of these steps in HIV-1 expression may be affected by CDK8/19 kinase inhibition, we assessed recruitment of RNA Pol II to the LTR using ChIP-qPCR. We found that treatment of unstimulated cells with Senexin A caused a significant decrease in association of RNA Pol II at the LTR RBE3 and RBE1 *cis*-elements that are located upstream of the enhancer region and near the transcriptional start site, respectively (Fig. 8*A*, compare Ve and SenA). Treatment with PEP005, a PKC agonist which imparts activation of NFκB, resulted in a significant increase in LTR associated RNAPII (Fig 8*A*, compare Ve and PEP005), but this effect is blocked by the CDK8/19 inhibitor Senexin A (Fig. 8*A*, compare PEP005 and PEP005/SenA). These results indicate that CDK8/19 kinase activity is necessary for efficient recruitment of RNA Pol II to the LTR.

**Figure 8.**
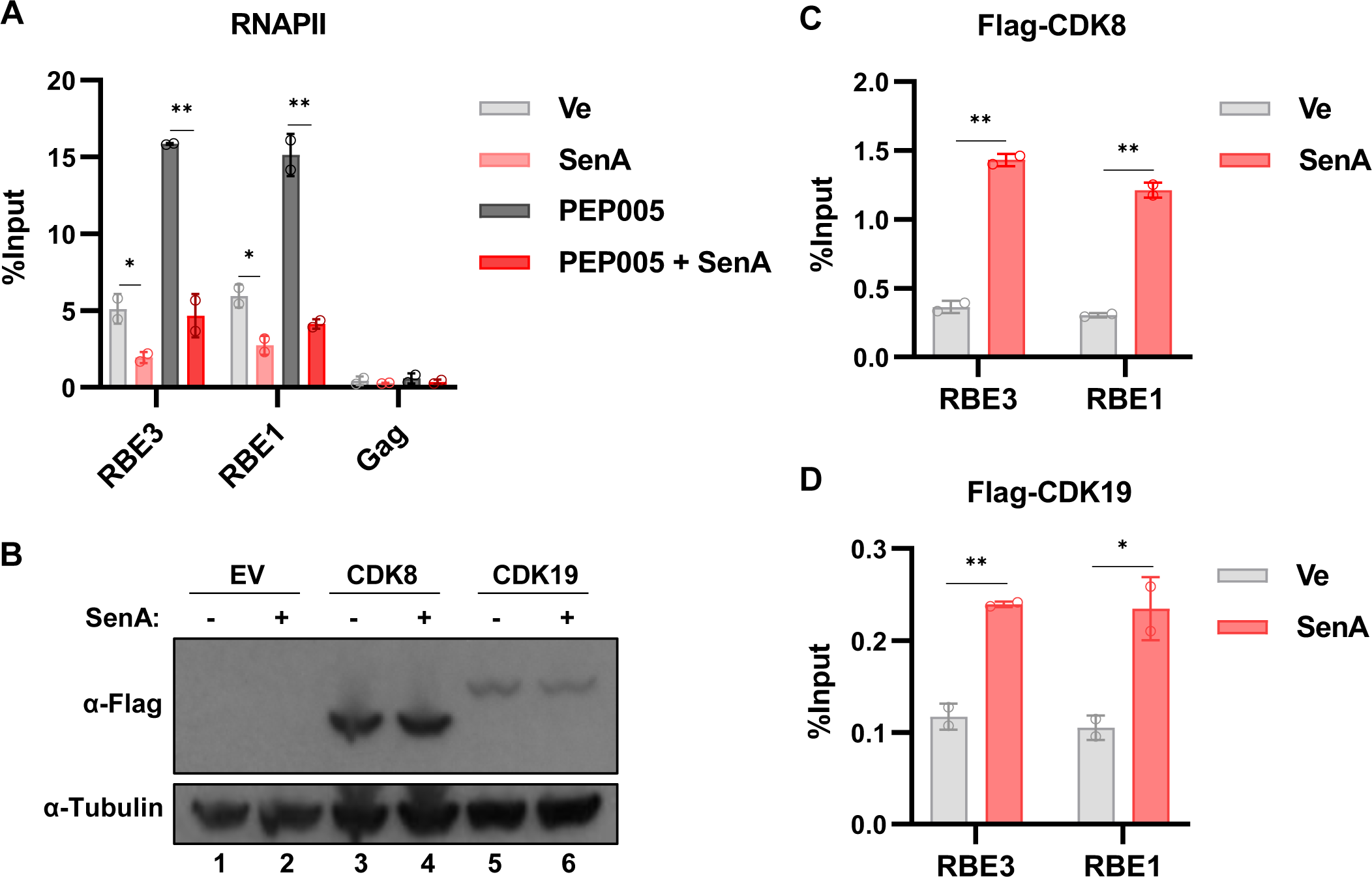
Inhibition of CDK8/19 impairs recruitment of RNA Polymerase II to the HIV-1 LTR. **A:** JLat10.6 cells were incubated for 20 hrs with a DMSO vehicle control (Ve), 10 μM Senexin A, 10 nM PEP005, or were pre-treated for 1 hr with 10 μM Senexin A, after which 10 nM PEP005 was added. ChIP was performed using an anti-RNAPII antibody and analyzed with qPCR with primers specific for the HIV-1 LTR RBE1 and RBE3 regions, or an intragenic Gag site. ChIP-qPCR results are normalized by subtraction of values produced with sample paired non-specific IgG immunoprecipitation (*n* = 2, mean ± SD). **B:** JLat10.6 cells were transduced with a non-expressing lentivirus (EV, lanes 1 and 2) or lentiviral vectors expressing Flag-tagged CDK8 (lanes 3 and 4) or Flag-tagged CDK19 (lanes 5 and 6). Transduced cells were left untreated (Sen A -) or incubated for 20 hrs with 10 μM Senexin A (Sen A +) after which cellular lysate was extracted and immunoblotted with antibodies against Flag and Tubulin. **C, D:** JLat10.6 cells expressing Flag-tagged CDK8 (C) or Flag-tagged CDK19 (D) were treated with DMSO vehicle control (Ve) or 10 μM Senexin A. Following 20 hrs incubation, ChIP was performed using anti-Flag antibody, and qPCR was used to detect enrichment at the RBE3 and RBE1 LTR sites. ChIP-qPCR results are normalized by subtraction of values produced with sample paired non-specific IgG immunoprecipitation (*n* = 2, mean ± SD).

Next, we examined the effect of CDK8/19 kinase inhibition on localization of CDK8 and CDK19 to the HIV-1 LTR. Treatment of cells expressing Flag-tagged CDK8 or CDK19 with Senexin A does not appear to affect abundance of these proteins (Fig. 8*B*), but interestingly, we observe increased association of both kinases with the LTR in cells treated with this inhibitor (Fig. 8*C*, 8*D*). It is possible that direct interaction of the CDK8/19 Kinase Module (CKM) negatively regulates HIV expression, which is consistent with a previous report indicating that CDK8 is evicted from the HIV-1 LTR upon T cell stimulation and induction of virus expression^57^. These observations would support a view that interaction of mediator kinase module with the promoter has a negative regulatory effect on transcription, a function that was also indicated from experiments with yeast^58^ and transcription reactions *in vitro*^59^. However, decreasing promoter transcriptional activity through TFIIH inhibition was found to enrich core promoter associated Mediator in yeast^58^. It is possible that suppressed LTR transcription in response to CDK8/19 inhibition causes reduced kinetics and allows our ChIP experiment to capture Flag tagged CDK8 or CDK19.

### CDK8 and CDK19 are activators of HIV-1 expression

The results above demonstrate that chemical inhibition of the CDK8/19 kinases inhibits HIV-1 basal and activated expression in Jurkat T cells and normal CD4^+^ lymphocytes. All current small molecule inhibitors of the mediator kinases do not differentiate between effects on CDK8 and CDK19^29 31^. Although these kinases are generally considered to have overlapping or redundant function for the mediator kinase module^60 31^, the role of CDK19 is less well understood and some evidence suggests specific functions for CDK19^61^. To assess the specific role of CDK8 in regulating HIV-1 expression, we used shRNA mediated knockdown of *CDK8* in both the mHIV-Luciferase (Fig. S3*A*) and JLat10.6 (Fig. S3*B*) cell lines. We found that knockdown of *CDK8* by shRNA in the mHIV-Luciferase cell rendered the integrated provirus significantly impaired for PMA induced luciferase reporter expression (Fig. S3*C*). Curiously however, we found that depletion of *CDK8* expression by shRNA in the JLat10.6 cell line did not inhibit induction of GFP upon PMA treatment (Fig. S3*D*). To examine the effect of CDK8 on HIV-1 transcription in more detail, we performed CRISPR-Cas9 gene editing to generate mHIV-Luciferase (Fig. 9*A*, lanes 2-7) and JLat10.6 (Fig. 9*A*, lanes 9-11) *CDK8* knockout (*CDK8* KO) cell lines. Consistent with results from shRNA knockdown, we found that all of the *CDK8* KO mHIV-Luciferase cell lines displayed a significant defect for reactivation of expression in response to PMA (Fig. 9*B*). However, similar to results with shRNA knockdown, we did not observe an effect of the *CDK8* KO in JLat10.6 cells on proviral expression in response to PMA (Fig. 9*C*). These observations indicate that CDK8 function is required for reporter gene expression in the mHIV-Luciferase but not in the JLat10.6 line. Considering that the chemical inhibitors of CDK8/19 inhibit PMA induced HIV-1 expression in both of these lines (Fig. 1), the differential effect of *CDK8* disruption in the mHIV-Luciferase and JLat10.6 lines may indicate functional differences between CDK8 and CDK19, a possibility we discuss further below. We do note that JLat10.6 cells typically express slightly higher levels of CDK8 protein than mHIV-Luciferase cells as determined by immunoblotting (Fig. 9*A,* Fig. S4).

**Figure 9.**
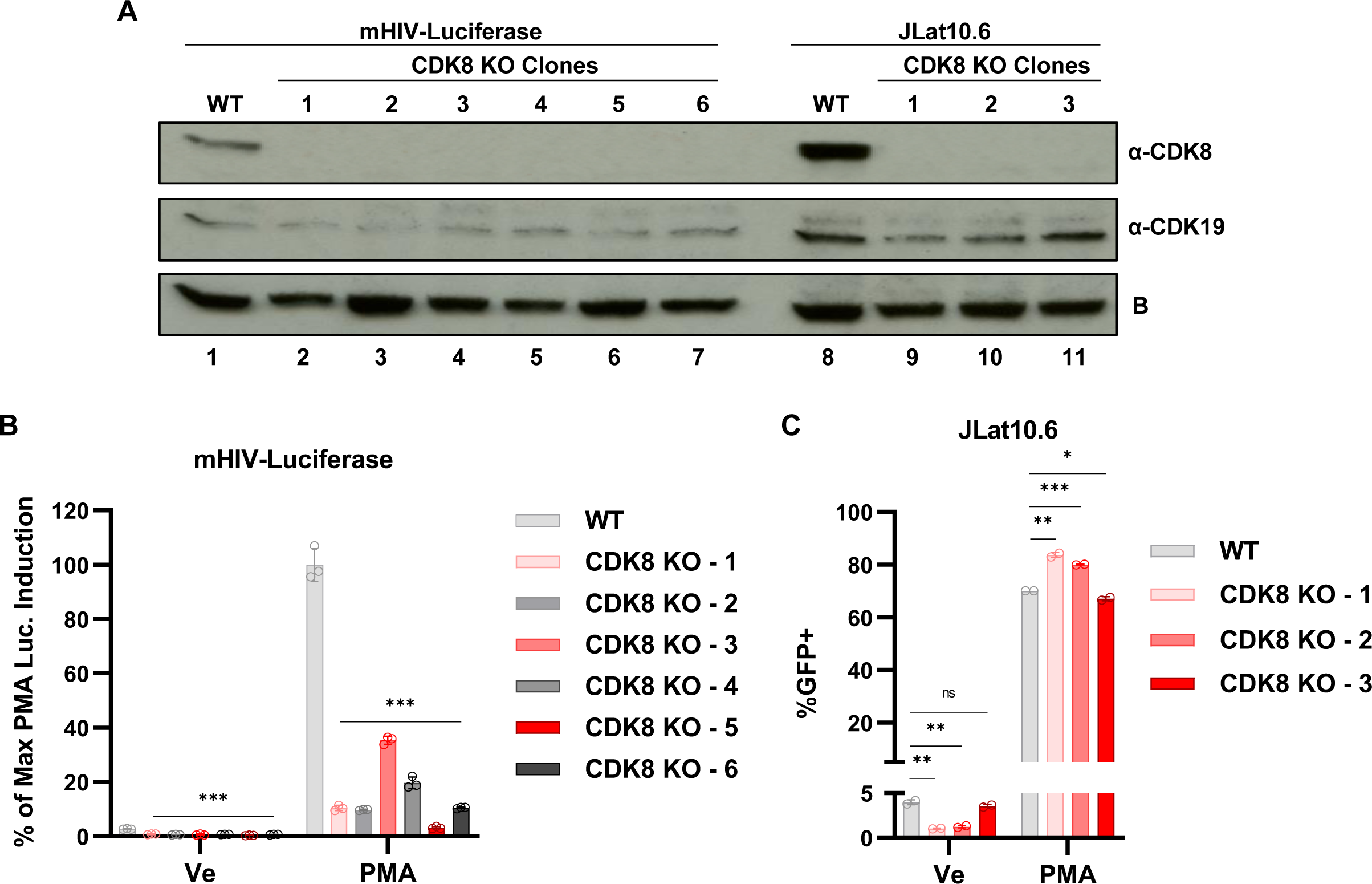
Effect of *CDK8* gene knockout on PMA induced HIV-1 expression. **A:** *CDK8* knockout clones were generated in the mHIV-Luciferase (lanes 2-7) and JLat10.6 (lanes 9-11) cell lines. Cell lysates from WT mHIV-luciferase (lane 1), WT JLat10.6 (lane 8), or the indicated KO clonal line were immunoblotted with antibodies against CDK8 and CDK19. (B) indicates a background signal that is produced by Rabbit antibodies against CDK19 that is unaffected by any condition we have examined. **B:** WT mHIV-luciferase or the *CDK8* KO clones (A) were left untreated (Ve, DMSO) or stimulated with 10 nM PMA for 4 hrs after which luciferase activity was measured (*n* = 3, mean ± SD). **C:** WT JLat10.6 cells or *CDK* KO clones (A) were left untreated (Un) or stimulated with 10 nM PMA for 20 hours after which the proportion of GFP expressing cells was measured by flow cytometry (*n* = 2, mean ± SD).

Previously, we observed differential expression of HIV-1 in response to PMA between the mHIV-Luciferase and JLat10.6 *CDK8* KO Jurkat cell lines (Fig. 9). These are two distinct clonal lines and as such, we sought to examine the effect of *CDK8* abrogation on expression of a heterogenous population of proviruses. For this analysis we used the replication incompetent Red-Blue-HIV-1 (RBH) dual HIV-1 reporter which expresses all viral gene products apart from *Nef* and *Env*, and where BFP is expressed from the 5’ LTR, and mCherry from an internal CMV promoter (Fig. 10*A*)^42^. Wildtype or *CDK8* KO Jurkat T cells were infected with RBH, and the populations were subsequently treated with a vehicle control (Ve) or an LRA 4 days post infection; flow cytometry was performed 1 day later to examine BFP and mCherry expression (Fig. 10*B,* Fig. S5). Importantly, we found that the proportion of productively infected cells in the otherwise untreated population was significantly lower in *CDK8* KO Jurkat cells relative to WT (Fig. 10*C*, Ve). The proportion of productively infected cells treated with the PKC agonists PMA and PEP005 was significantly increased, but interestingly, this proportion was nearly identical in the *CDK8* KO line (Fig. 10*C*, PMA, PEP005). In contrast we found that treatment of RBH infected *CDK8* KO cells with the histone deacetylase inhibitor SAHA, or the bromodomain inhibitors JQ1 and IACS-957^18^ failed to recapitulate the level of productive infections observed for wildtype cells (Fig. 10*C*, SAHA, JQ1, IACS-9571).

**Figure 10.**
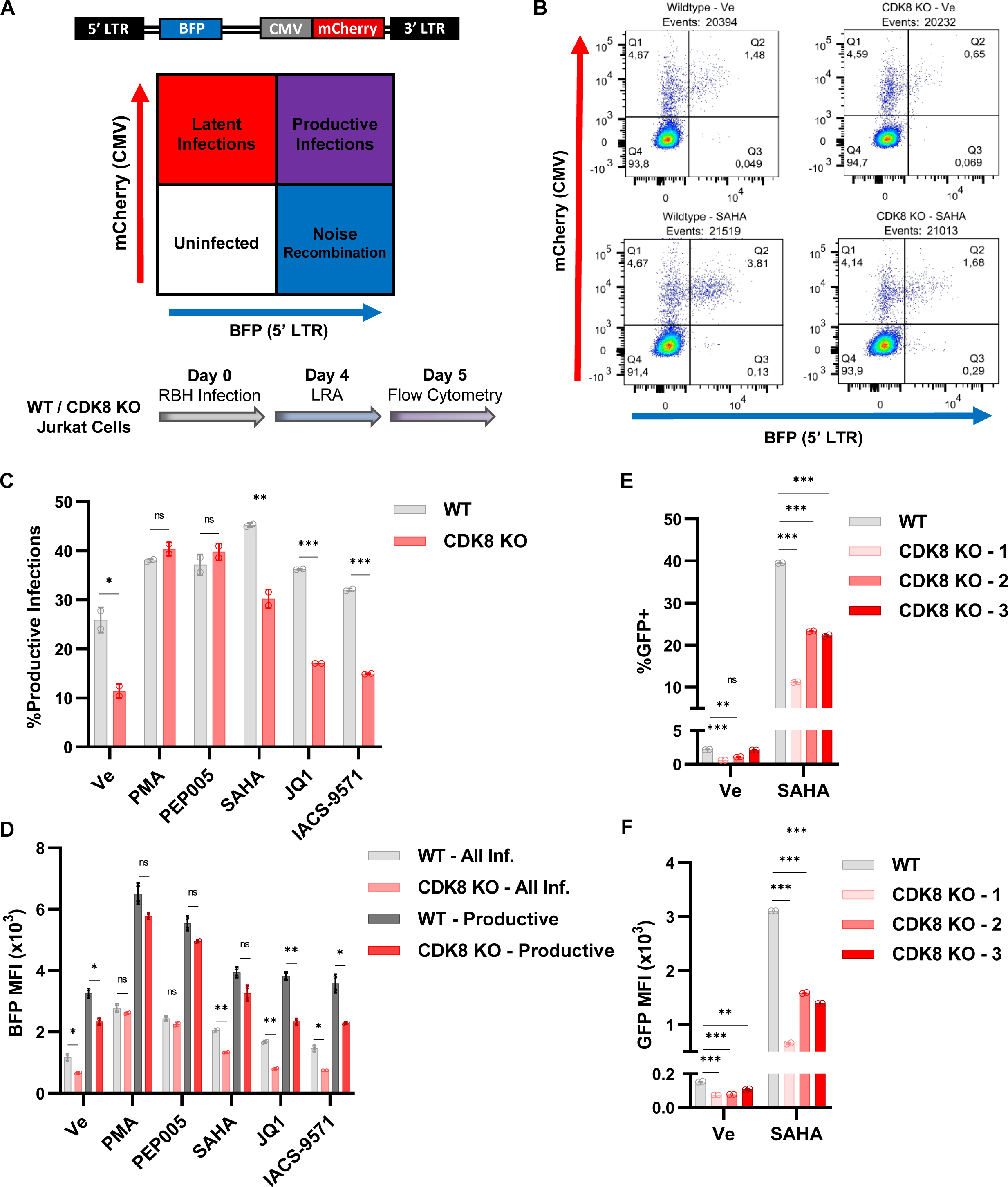
*CDK8* knockout desensitizes HIV-1 to several latency reversal agents. **A:** Schematic description of the Red-Blue-HIV-1 (RBH) dual reporter assay. RBH infection produces a distinct scatter plot upon flow cytometric analysis that is similar to RGH (Fig. 4A), but with BFP expression being driven by the 5’ LTR instead of GFP ^42^. We first infected wildtype or *CDK8* KO Jurkat T cells with RBH. 4 days post-infection, we treated the cells with a latency reversal agent and assessed HIV-1 expression following 20 hrs of incubation. **B:** Representative flow cytometric scatter plots of wildtype (left) or *CDK8* KO (right) Jurkat cells following incubation with DMSO (Ve) (top) or SAHA (bottom). **C:** Wildtype or *CDK8* KO cells were treated with DMSO vehicle control (Ve), 10 nM PMA, 10 nM PEP005, 10 μM SAHA, 10 μM JQ1, or 10 μM IACS-9571, and flow cytometry was performed 20 hrs later to determine the percentage of productive infections (*n* = 2, mean ± SD). **D:** As in (C), but the Mean Fluorescence Intensity (MFI) of LTR driven BFP expression is depicted. All Infections measures the expression of BFP in the populations of cells that are mCherry+/BFP+ and mCherry+/BFP+ while Productive measures BFP expression in population that is mCherry+/BFP+ (*n* = 2, mean ± SD). **E**: WT or *CDK8* KO JLat10.6 cells (A) were left untreated or incubated with 10 μM SAHA; 20 hours later flow cytometry was performed to determine the proportion of GFP+ cells (*n* = 2, mean ± SD). **H:** As in (G), but viral expression is reported as the Mean Fluorescence Intensity (MFI) of GFP expression (*n* = 2, mean ± SD).

Beyond assessing the ratio of productive versus latent infections, we took a closer examination of the effect of *CDK8* KO on LTR driven BFP expression. This analysis was performed by independently evaluating the BFP mean fluorescence intensity of all infected cells (mCherry+/BFP-and mCherry+/BFP+, All Inf.) and the productive proviral infections (mCherry+/BFP+, Productive). Comparison of BFP expression of all infections showed that *CDK8* KO cells were associated with decreased LTR activity upon treatment with a vehicle control (Ve), SAHA, JQ1, or IACS-9571 but no difference was apparent upon stimulation with PMA or PEP005 (Fig. 10*D*, All Inf.). This is expected as we are examining BFP expression for the whole of integrated proviruses and the results mirror the effect of *CDK8* KO on the productive infection ratio for the indicated treatment (Fig. 10*C*). Next, we examined LTR driven BFP expression of the proviruses that established productive infections (the mCherry+/BFP+ population). Here, we observed a slight but non-significant decrease in viral transcription of *CDK8* KO cells following treatment with PMA, PEP005, or SAHA (Fig. 10*D*). However, the productive proviral infections of *CDK8* KO cells following incubation with a vehicle control, JQ1, or IACS-9571 displayed greatly reduced LTR expression as compared to wildtype (Fig. 10*D*, Productive). Collectively, these results display that CDK8 regulates basal HIV-1 transcription as well as activation in response SAHA, JQ1, and IACS-9571. However, CDK8 is not required for viral induction in response to PMA or PEP005. As the CDK8/19 dual inhibitors reduce HIV-1 expression in response to these agonists (Fig. 1, 3), CDK19 must be sufficient to mediate LTR induction in response to PKC activation as mediated by PMA or PEP005.

The RBH reporter virus allowed us to assess a heterogenous population of proviruses and identify latency reversal pathways that are strongly or weakly dependent on CDK8. For instance, CDK8 was not required for viral activation as induced by the PKC agonists PMA or PEP005 but was necessary for LTR responsiveness to the histone deacetylase inhibitor SAHA, the BRD4 bromodomain inhibitor JQ1, and the TRIM24 bromodomain inhibitor IACS-9571 (Fig. 10*C*, 10*D*). To characterize this further, we examined the reactivation of wildtype and *CDK8* KO JLat10.6 cells in response to the HDACi, SAHA. Although JLat10.6 cells did not require *CDK8* for PMA mediated reactivation (Fig. 9*C*), we found that *CDK8* null cells displayed substantially dampened responsiveness to SAHA (Fig. 10*E*, 10*F*). Altogether, these results indicate that CDK8 and CDK19 may have overlapping functions for viral response to PMA or PEP005 but divergent function for reactivation by the HDAC inhibitor SAHA.

### CDK8/19 inhibitors suppress HIV-1 reactivation in CD4^+^ T cells

Because inhibition of CDK8/19 impairs HIV-1 expression in multiple models of latency and normal CD4^+^ T cells, we wondered if these inhibitors could also block reactivation of virus in primary CD4^+^ T cells from individuals with HIV-1 receiving suppressive antiretroviral therapy (ART). Treatment of CD4^+^ T cells from five participants with a combination PMA and ionomycin stimulated HIV-1 mRNA expression in all cases (Fig. 11*A*, PMA/Ion). However, Senexin A inhibition of CDK8/19 kinase activity robustly limited induction of HIV-1 viral expression by PMA and ionomycin in CD4^+^ T cells from all ART-treated participants (Fig. 11*A*, PMA/Ion/SenA). We observe a similar effect on expression of *IL2* in these samples (Fig. 11*B*), which is consistent with *IL2* and HIV-1 both being regulated by T cell signaling ^13^. Senexin A caused minor alterations, but more divergent effects on expression of *CD69*, a marker of T cell activation in these treatments (Fig. 11*C*). Importantly, Senexin A treatment had no discernable effect on cell viability at concentrations where we observe inhibition of HIV-1 provirus reactivation (Fig. 11*D*). Collectively, these observations indicate that chemical inhibitors of CDK8/19, including Senexin A, may represent potential therapies to suppress HIV-1 provirus for elimination of latently infected cells in individuals on ART.

**Figure 11.**
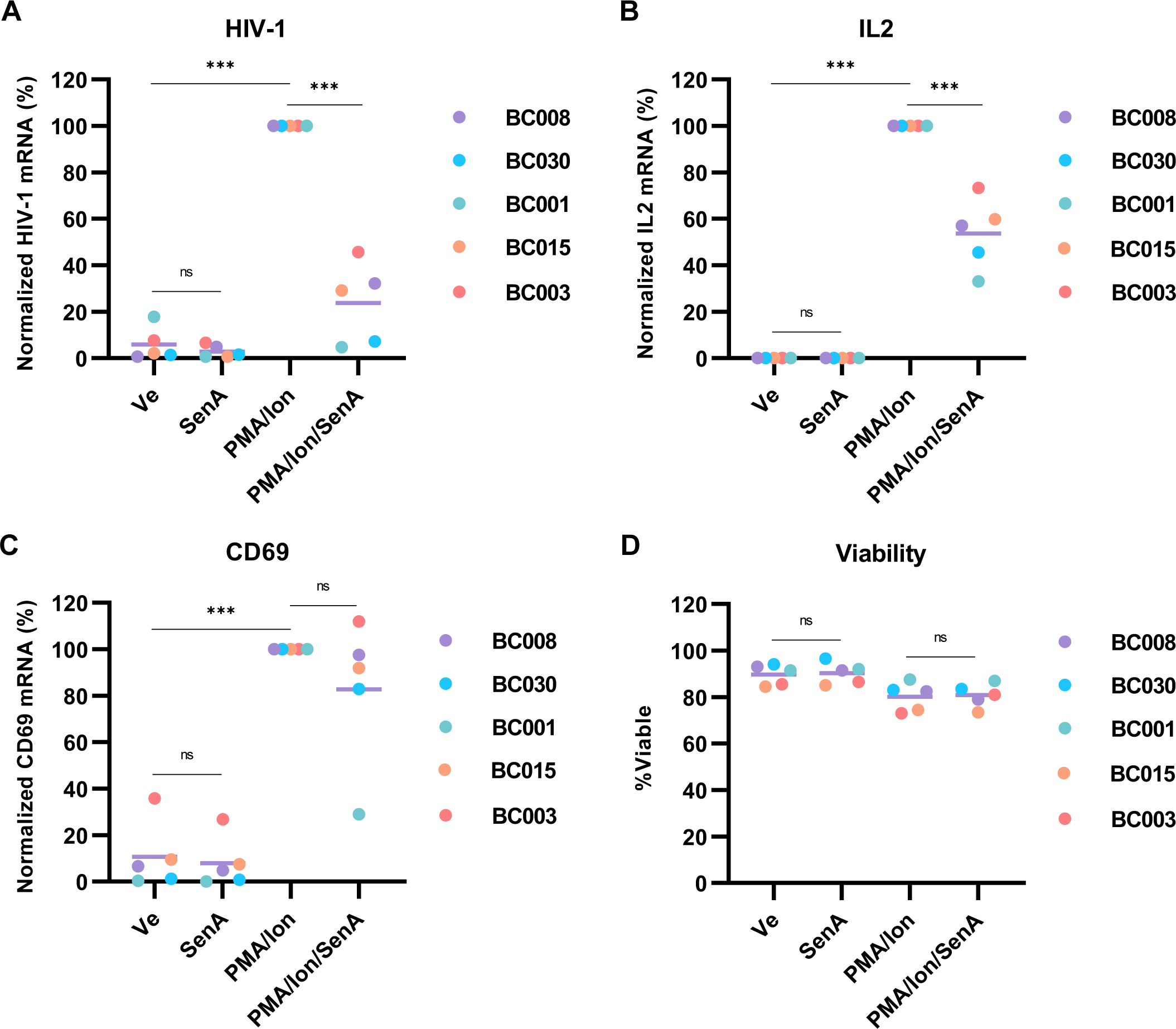
Senexin A inhibits HIV-1 provirus reactivation in PBMCs from patients on ART. **A-C:** CD4^+^ PBMCs, isolated from HIV-1 infected individuals on ART, were treated for 20 hrs with DMSO (Ve), 10 μM Senexin A (Sen A), 10nM PMA and 1μM Ionomycin (PMA/ Ion), or were pre-treated with 10 μM Senexin A for 1 hr prior to the addition of 10 nM PMA/ 1μM Ionomycin. Intracellular RNA was extracted and analyzed using RT-PCR with oligos specific for multiply spliced Tat-Rev HIV-1 mRNA transcripts (A), *IL2* (B), and *CD69* (C). Expression of the indicated mRNA is normalized to *GAPDH* (per indicated patient ID, the mean of *n* = 2 – 3 is shown). **D**: Viability of cells following treatment as in (A – C) was determined (per indicated patient ID, the mean of *n* = 2 is shown).

## Discussion

Expression of chromosomally integrated HIV-1 provirus in T cells is dependent upon cellular factors that bind the LTR enhancer region and are regulated by signaling pathways stimulated by T cell receptor engagement^62 14^. Of central importance to this response is NF-κB, which is activated by phosphorylation and inhibition of its cytoplasmic inhibitor I-κB, allowing translocation of NF-κB p65 RelA to the nucleus where it binds two sites within the LTR enhancer region and activates transcription^63 64^. Previous work showed that CDK8/19 is co-recruited with NF-κB to promoters, and that CDK8/19 inhibitors impair transcriptional activation by NF-κB^24^. Additional transcriptional activators regulated by CDK8 also bind the 5’ HIV-1 LTR including, β-catenin TCF/LEF^23^ and STAT1/3^25^, which is consistent with our findings that CDK8/19 inhibitors, *CDK8* knockdowns, and *CDK8* gene knockouts inhibit reactivation of HIV-1. We find that CDK8/19 inhibitors impair induction of HIV-1 provirus in cell line models and newly infected CD4^+^ T cells, and in response to a variety of latency reversing agents, including PEP005, SAHA and JQ1. Importantly, we find that the CDK8/19 inhibitor Senexin A prevents reactivation of latent virus expression in CD4^+^ T cells from individuals with HIV receiving suppressive antiretroviral therapy. These results indicate that chemical inhibitors of CDK8/19, some of which are presently in clinical trials for various cancers^65^, could be useful to provide longstanding and durable suppression of the latent HIV-1 reservoir.

We find that Senexin A impairs recruitment of RNA Polymerase II to the LTR promoter, which suggests that it causes a defect in transcriptional activation by LTR-bound transcription factors. Treatment of T cells with the PKC agonist PEP005 causes significantly enhanced association of RNA Pol II with the LTR, which likely represents the effect of LTR-associated NF-κB p65 and recruitment of general transcription factor complexes, including mediator and P-TEFb^66 67^. Stimulation of NF-κB p65 by TNF-α was shown to cause recruitment of TFII-H/ CDK7, which was identified as a rate limiting event for reactivation of provirus from latency^57^. We find that Senexin A inhibits association of RNA Pol II to the LTR in untreated cells and those stimulated with the PKC agonist PEP005 (Fig. 8*A*). Latent HIV-1 provirus is associated with RNA Pol II that is paused immediately downstream of the transcriptional start site, following synthesis of the nascent TAR RNA ^68^, and activation of HIV-1 involves stimulation of elongation from the viral promoter through recruitment of P-TEFb containing CDK9 and cyclin T1^69^. However, the factors and mechanisms involved in establishing paused RNA Polymerase and associated GTFs at the latent HIV-1 core promoter have not been determined. Our results indicate that CDK8 activity might be required for establishment of paused RNA Pol II at the promoter. Consistent with this possibility the mediator kinase module, including CDK8 was found associated with the latent virus promoter^57^.

The CDK8/19 inhibitors Senexin A and BRD6989 have similar effects on CDK8 and CDK19 *in vitro*, and presumably do not discriminate for inhibition of these paralogs *in vivo*^70 35^. Consequently, it is interesting that these compounds cause only partial inhibition of HIV-1 reactivation in cell line models, but that knockout of the *CDK8* gene in the mHIV-luciferase line nearly completely inhibits reactivation in most knockout clones (Fig. 9*B*). This observation may indicate that CDK8 has a structural role for regulation of HIV-1 transcription in addition to catalytic function. Furthermore, although depletion of *CDK8* by shRNA mediated knockdown or genetic ablation caused significant inhibition of HIV-1 expression in the mHIV-Luciferase line, they have no effect on PMA or PEP005 stimulated GFP reporter expression in the JLat10.6 line. Potential mechanistic differences between these kinases might depend upon specific properties of the JLat10.6 and mHIV-Luciferase cell lines, for example differences in the site of chromosomal integration. Also, we note that luciferase is expressed in the mHIV-Luciferase as a fusion with Gag from an unspliced transcript, whereas expression of GFP in JLat10.6 is dependent upon production of a spliced sub-genomic RNA. Spliced HIV-1 transcripts, including that encoding Nef, which is replaced by GFP in the JLat10.6 line^71^, are expressed early in infection or reactivation from provirus latency. Splicing of sub-genomic transcripts is suppressed later in infection by the viral Rev protein, which enables synthesis of full length genomic RNAs^72^. In contrast, expression of luciferase as a fusion with Gag in the mHIV-Luciferase cell line does not require mRNA splicing (Fig. 1*B*)^73^. Alternatively, the differential effect of CDK8 and CDK19 between reactivation of HIV-1 in these lines may be dependent upon expression of additional HIV-1 gene products, as the JLat10.6 line is capable of expressing all viral proteins, apart from Nef ^1^, whereas coding sequences for all viral accessory factors is deleted in the mHIV-luciferase line^73^.

Additionally, there was an apparent overlap in phenotype between C*DK8* KO cells and CDK8/CDK19 inhibited cells when we examined the heterogenous population of proviruses that was generated by RBH infection. In response to SAHA, JQ1, and IACS-9571, *CDK8* ablation mimicked the effect of CDK8/19 inhibition, causing suppressed LTR transcriptional activity. However, unlike CDK8/19 inhibition, loss of CDK8 had no effect on proviral induction in response to the PKC agonists PMA or PEP005. Furthermore, activation of HIV-1 expression in *CDK8* null JLat10.6 cell lines shared a similar pattern to RBH proviruses. Given these observations, it is likely that HIV-1 reactivation in response to SAHA, JQ1, and IACS-9571 is largely dependent upon CDK8, while CDK19 is generally sufficient for PKC mediated transcriptional induction. The mechanism(s) contributing to these differences are not understood, but likely represent distinct functions for CDK8 and CDK19, as Senexin A and BRD6989 inhibit HIV-1 expression in response to all of these stimuli.

The mediator kinases were previously shown to have distinct functions, where it was shown that CDK19 but not CDK8 has a significant role for regulation of cell cycle progression of mouse hematopoietic stem cells^61^. We also observed that Senexin A and BRD6989 did not have an effect on HIV-1 expression in HEK293T or HeLa cells, indicating possible cell-type-specific functions for the mediator kinase. This latter result could account for a previous report indicating that the mediator kinases do not affect HIV-1 expression^74^. Overall however, a more detailed understanding of the mechanistic role of CDK8 and CDK19 for regulation of transcription will be required to resolve these discrepancies.

Latent HIV-1 proviruses are known to produce sporadic transcripts through transcriptional noise^75 76^, such that even the most strongly repressed provirus will occasionally produce viral transcripts^44 4^. Considering these observations, it is possible that stochastic expression of latent provirus may contribute to maintenance of the latent population in tissue compartments where ART may not be capable of preventing local spread of virus^4^. Consequently, one proposed therapeutic strategy, designated block and lock, would involve intervention to suppress stochastic expression of provirus in patients on ART to promote deep latency, which may enable elimination of the latently infected population by cell lifespan decay^86^. The results presented here indicate that inhibitors of the CDK8/19 mediator kinases may be an important contribution towards this strategy.

## Materials and Methods

### Cell culture, virus culture, and lentiviral transduction

Jurkat E6-1, SupT1, Jurkat Tat mHIV-Luciferase, JLat10.6, ACH2, and U1 cells were cultured in Roswell Park Memorial Institute (RPMI) 1640 medium supplemented with 10% FBS, penicillin [100 units/ml], streptomycin [100 g/ml], and L-glutamine [2 mM]. HEK293T and TZM-bl cells were cultured in Dulbecco’s modified Eagle’s (DMEM) medium supplemented with 10% fetal bovine serum (FBS), penicillin [100 units/ml], streptomycin [100 g/ml], and L-glutamine [2 mM]). All cell lines were incubated in a humidified 37°C and 5% CO_2_ atmosphere.

Human Peripheral Blood CD4^+^ T cells, purchased from STEMCELL Technologies (Catalog # 200-0165), were cultured in RPMI supplemented with 10% FBS, penicillin (100 U/mL), streptomycin (100 mg/mL), and 30 U/mL IL2. RGH infection was performed as previously described^45^. Briefly, cells were incubated in the presence of Dynabeads^TM^ Human T-Activator CD3/CD28 beads for three days. Subsequently, the beads were removed, and cells infected with RGH at a multiplicity of infection (M.O.I.) causing less than 5% of the population to be infected.

Peripheral Blood Mononuclear Cells (PBMC) from participants with HIV-1 on ART were isolated from whole blood by density gradient centrifugation using Lymphoprep^TM^ and SepMate^TM^ tubes (StemCell Technologies), and cryopreserved. Upon thawing, PBMCs were cultured in RPMI supplemented with 10% FBS, penicillin [100 units/ml], streptomycin [100 g/ml], and L-glutamine [2 mM]. Samples from participants were collected with written informed consent under a protocol jointly approved by the research ethics boards at Providence Health Care/UBC and Simon Fraser University (certificate H16-02474).

Vesicular stomatitis virus G (VSV-G) pseudotyped viral stocks were produced by co-transfecting HEK293T cells with a combination of viral molecular clone, psPAX, and VSV-G at a ratio of 8 μg: 4 μg: 2 μg. Transfections were performed with polyethylenimine (PEI) at a ratio of 6:1 (PEI:DNA) in Gibco Opti-MEM^tm^. Lentiviral infections were performed by plating 1×10^6^ cells in 24-well plates with media containing 8 μg/mL polybrene and the amount of viral supernatant to give the desired multiplicity of infection (M.O.I.) as indicated. Plates were subsequently spinoculated for 1.5 hrs at 1500 rpm.

### shRNA knockdown

JLat10.6 or Jurkat mHIV-Luciferase cells were transduced with a pLKO empty vector or pLKO shRNA expressing lentivirus at a M.O.I. ∼10. Infected JLat10.6 cells were selected by culturing in 1 μg/mL puromycin, while mHIV-Luciferase cells had media supplemented with 3 μg/mL puromycin. All experiments were performed 3 – 10 days following shRNA transduction. MISSION shRNA clones (Sigma) used for knockdown are as follows: CDK8-1, TRCN0000000489 – ATGTCCAGTAGCCAAGTTCCA (3’ UTR); CDK8-2, TRCN0000350308 – CTAACGTCAGAACCAATATTT (CDS); CDK8-3, TRCN0000350344 – GCTTACCATGGACCCAATAAA (CDS).

### CDK8 knockout

CRISPR-Cas9 was used to generate clonal CDK8 knockout cells from mHIV-Luciferase and JLat10.6 cells. Cas9 (pU6_CBh-Cas9-T2A-BFP: Addgene #64323) and gRNA cassette (pSPgRNA: Addgene #47108) plasmids had sequences that target genomic *CDK8*. These plasmids were than co-transfected into 2×10^6^ cells using the Neon Transfection System (Invitrogen) as per the manufacturer’s instructions with the following settings: Voltage, 1350 V; Width, 20 ms; Pulse number, 3x. Knockout cells were isolated by live sorting (Astrios Flow Cytometer) BFP positive cells into 96-well plates containing complete RPMI 1640. Clones were expanded, and *CDK8* KO was validated by PCR genotyping and western blotting. *CDK8* gRNA target sequences were GGCCTCAGAGGCTGTGACAA and GTCTGATGTGAGTACTGTGG.

### Immunoblotting

Western blotting was performed as previously described (Hashemi *et al*, 2018). Antibodies were as follows: Tubulin - Abcam ab7291 (1:20000 – 1:40000), CDK8 - Abcam ab229192 (1:4000 – 1:10000), CDK19 – Sigma-Aldrich SAB4301196 (1:750), Goat Anti-Rabbit-HRP – Abcam ab6721 (1:2000000), Goat Anti-Mouse-HRP – Pierce 1858413 (1:20000).

### Chromatin Immunoprecipitation

ChIP-qPCR was performed as previously described (Horvath *et al*, 2023) (Horvath *et al*, 2023). The following antibodies were used for IP: RNAPII - Abcam ab26721 (5 μg), Flag - Sigma Aldrich F3165 (5 μg). Cycling parameters were as follows: 50 °C, 2 min, 1x; 95 °C, 10 min, 1x; 95 °C, 15 sec, 60 °C, 1 min, 40x. Oligos used for qPCR are as follows: RBE3, Fwd 5’ AGCCGCCTAGCATTTCATC (nucleotide coordinates 270-288 in the HIV-1 subtype B reference strain HXB2), Rev 5’ CAGCGGAAAGTCCCTTGTAG (HXB2 nucleotides 363-344); RBE1, Fwd 5’ AGTGGCGAGCCCTCAGAT, Rev 5’ AGAGCTCCCAGGCTCAGATC; Gag: Fwd 5’ AGCAGCCATGCAAATGTTA, Rev 5’ AGAGAACCAAGGGGAAGTGA.

### Luciferase reporter assays

For TZM-bl cells, 2×10^4^ cells were plated with 100 μL DMEM per well in 96-well plates. Following 24 hrs, cells were incubated with the indicated concentration of drug for 4 hrs and luciferase expression was measured. For Jurkat mHIV-Luciferase cells, 1×10^5^ luciferase expressing cells were plated with 100 µL media in 96-well plates. Luciferase activity was measured after the indicated time of treatment. Measurements were performed using the Superlight^TM^ luciferase reporter Gene Assay Kit (BioAssay Systems) as per the manufacturer’s instructions and 96-well plates were read in a VictorTM X3 Multilabel Plate Reader.

### RT-PCR

RNA was extracted from cells using the RNeasy Kit (Qiagen) and analyzed with the Quant Studio 3 Real-Time PCR system (Applied Biosystems) using *Power* SYBR® Green RNA-to-CT™ 1-Step Kit (Thermo Fisher) as per the manufacturer’s instructions. RT-PCR data was normalized to GAPDH expression using the ΔΔCt method as previously described^77^. Cycling parameters were as follows: 48 °C, 30 min, 1x; 95 °C, 10 min, 1x; 95 °C, 15 sec, 60 °C, 1 min, 60x. Primers were as follows: IL2, Fwd 5’ AACTCACCAGGATGCTCACA, Rev 5’ GCACTTCCTCCAGAGGTTTGA; CD69, Fwd 5’ TCTTTGCATCCGGAGAGTGGA, Rev 5’ ATTACAGCACACAGGACAGGA; HIV-1 mRNA (multiply spliced Tat-Rev transcripts), Fwd 5’ CTTAGGCATCTCCTATGGCAGGA, Rev 5’ GGATCTGTCTCTGTCTCTCTCTCCACC; GAPDH, Fwd 5’ TGCACCACCAACTGCTTAGC, Rev 5’ GGCATGGACTGTGGTCATGAG.

### Flow cytometry

Cells were treated as indicated in the figure legends. For flow cytometric analysis, human T cell lineages were suspended in PBS while HEK293T cells were suspended in PBS containing 10% trypsin-EDTA to prevent aggregation. A BD Biosciences LSRII-561 system was used for flow cytometry with threshold forward scatter (FSC) and side scatter (SSC) parameters being set so that a homogenous population of live cells was counted (Fig. S6). FlowJo software (TreeStar) was used to analyze data and determine the indicated Mean Fluorescence Intensity (MFI).

### Statistics and reproducibility

All replicates are independent biological replicates and are presented as mean values with ± standard deviation shown by error bars. The number of times that an experiment was performed is indicated in the figure legends. P-values were determined by performing unpaired samples *t*-test with the use of GraphPad Prism 9.0.0. Statistical significance is indicated at \**P* < 0.05, \*\**P* < 0.01, or \*\*\**P* < 0.001, with n.s. denoting non-significant *P* ≥ 0.05.

## Data availability

All data supporting the findings of this study are available within the article or from the corresponding author upon reasonable request (I. Sadowski). RNA-seq data was obtained from NCBI GEO, accession GSE221851^78^.

## Acknowledgments

We thank Andy Johnson and Justin Wong of the UBC Flow Cytometry Facility for performing FACS analysis as well as for assistance with flow cytometry. We thank the laboratory staff at the BC Centre for Excellence in HIV/AIDS for processing PBMCs from study participants. This research was supported by program project grant F16-01210, from the Canadian Institutes of Health Research (CIHR), and F19-05392 Discovery Grant from the Natural Sciences and Engineering Research Council of Canada (NSERC) (to I.S.). PBMC collection from participants with HIV-1 was supported by CIHR project grant PJT-159625 (to ZLB). ZLB is supported by a Michael Smith Health Research BC Scholar Award. Maria Aristizabal and colleagues provided valuable comments on the manuscript.

## Author Contributions

Horvath R. M. performed all experiments. Brumme Z.L. provided PBMC samples from individuals on ART. Sadowski I. and Horvath R. M. wrote the paper.

